# Cryo-EM structure of the type III-E CRISPR-Cas effector gRAMP in complex with TPR-CHAT

**DOI:** 10.1101/2022.09.07.506877

**Authors:** Shuo Wang, Minghui Guo, Yuwei Zhu, Zhiying Lin, Zhiwei Huang

**Author notes:** These authors contributed equally: Shuo Wang, Minghui Guo, Yuwei Zhu.

## Abstract

The type III-E CRISPR-Cas effector protein, named gRAMP, is the largest single-unit CRISPR-Cas effector. A caspase-like peptidase (TPR-CHAT) gene often co-occurs with gRAMP gene clusters. However, the exact mechanism of the recognition and cleavage of target RNA of Sb-gRAMP, as well as the molecular architecture of the CRISPR-guided caspase complex, remains unclear. Here, we report the cryo-EM structures of the type III-E effector Sb-gRAMP-crRNA in a complex with TPR-CHAT, with and without target ssRNA at 3.0 and 2.9 Å, respectively. The overall structure of the gRAMP-crRNA-ssRNA-TPR-CHAT complex adopts an “L”-shaped conformation, consisting of a copy of gRAMP, a copy of TPR-CHAT, a 37-nt crRNA, and an 18-nt target ssRNA. The data presented in this manuscript reveal the mechanism of recognition of crRNA and target ssRNA by gRAMP, also, this work reports the structure of the CRISPR type III-E effector in complex with the binding partner TPR-CHAT, which provides vital clues for elucidating the functional relation between the CRISPR-Cas system and caspase peptidase.

## Introduction

Prokaryotes and viruses have been engaged in an evolutionary struggle for billions of years^1^. Bacteria and archaea employ clustered regularly interspaced short palindromic repeats (CRISPR)-Cas adaptive immune systems to protect against viral infection^2^. The CRISPR-Cas locus consists of two parts: CRISPR and CRISPR associated (Cas) genes. According to the latest phylogenetic classification, the CRISPR-Cas system can be divided into two classes, which can be further subdivided into six types^3^. Class 1 CRISPR-Cas systems are characterized by a multiprotein effector, and consist of type I, III, and IV CRISPR-Cas systems. Class 2 CRISPR-Cas systems feature a single nuclease protein, and consist of type II, V, and VI CRISPR-Cas systems. The type II Cas9 and type V Cas12 systems have been engineered as genome-editing tools, and they have been successfully applied in a broad range of organisms^4,5^. Compared with Cas9, the type VI Cas13 system exhibits RNA-guided RNase activity and the property of nonspecific cutting^6^. Recently, a new subtype of CRISPR-Cas, type III-E^3^, has been identified and named gRAMP^7^ or Cas7-11^8^. Unlike the traditional effectors of the type III CRISPR-Cas system, gRAMP is a single effector protein composed of four Cas7 proteins and one Cas11 protein fused together. Given its unique architecture and specific cleavage activity, gRAMP is expected to be engineered as a powerful RNA editing tool^8^. Interestingly, a gene encoding the caspase-like peptidase TPR-CHAT often co-occurs with gRAMP gene clusters^3^. TPR-CHAT interacts directly with gRAMP^7^, indicating a functional relation between the CRISPR-Cas system and caspase peptidase. However, the exact mechanism of the recognition and cleavage of target RNA of gRAMP from *Candidatus* “Scalindua brodae” (*Sb*-gRAMP)^7^, as well as the molecular architecture of the CRISPR-guided caspase complex, remains unclear. This study determined the cryo-EM (cryo-electron microscopy) structures of the type III-E effector *Sb*-gRAMP-crRNA in a complex with TPR-CHAT, with and without target ssRNA at 3.0 and 2.9 Å, respectively (Figure S1-2). The resulting maps were resolved well enough to facilitate the construction of a de novo model for the gRAMP-crRNA-ssRNA-TPR-CHAT complex (Figure 1A-B,S3).

**Figure 1.**
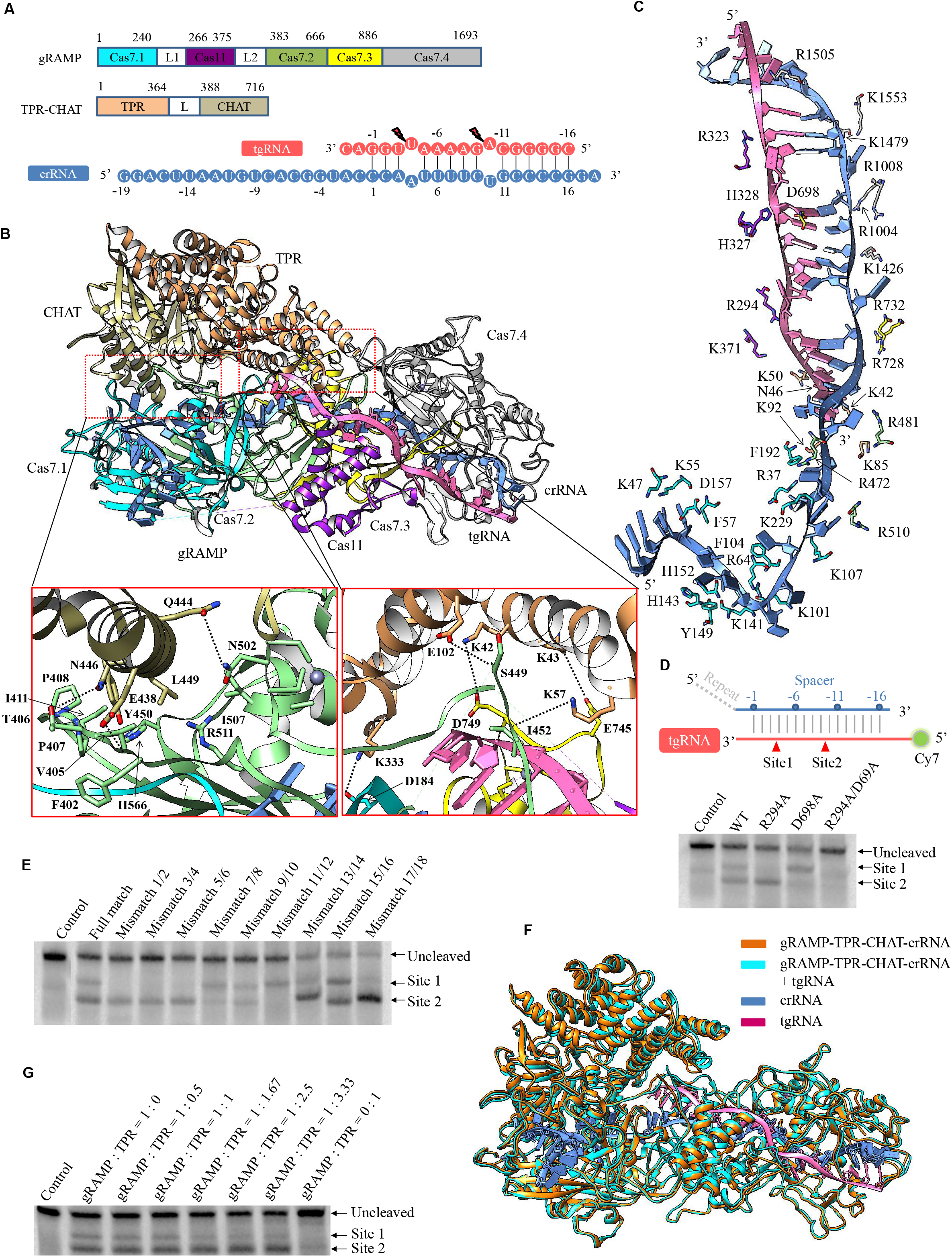
Cryo-EM structure of the gRAMP in complex with TPR-CHAT. **(A)** Graphic representation of the domain organization of the gRAMP and TPR-CHAT proteins. Schematic representation of the crRNA: target ssRNA heteroduplex. The target RNA cleavage sites are labeled. **(B)** Illustration of the overall structure of the gRAMP-crRNA-ssRNA-TPR-CHAT complex. The Cas7.1, Cas11, Cas7.2, Cas7.3, Cas7.4, TPR, and CHAT domains are shown in cyan, purple, green, yellow, gray, goldenrod, and sandy brown, respectively. **(C)** Illustration of the interactions between gRAMP and the bound nucleic acids. Colors used to represent interaction residues are the same as those shown in (**B**). **(D)** Cleavage activity analysis of WT gRAMP and its mutants. **(E)** In vitro target ssRNA cleavage by the WT gRAMP using mismatched RNAs. **(F)** Structural comparison of gRAMP-crRNA-TPR-CHAT with or without target ssRNA. The gRAMP-crRNA-TPR-CHAT complex is shown in cyan, and gRAMP-crRNA-ssRNA-TPR-CHAT is shown in orange. **(G)** In vitro target ssRNA cleavage assay of WT gRAMP in the presence of TPR-CHAT.

## Results

The overall structure of the gRAMP-crRNA-ssRNA-TPR-CHAT complex adopts an “L”-shaped conformation, consisting of a copy of gRAMP, a copy of TPR-CHAT, a 37-nt crRNA, and an 18-nt target ssRNA (Figure 1B). The length and the width of the gRAMP-crRNA-ssRNA-TPR-CHAT is approximately 145 Å and 96 Å, respectively. The gRAMP is composed of one Cas11 domain and four Cas7 domains. Four Cas7 domains (Cas7.1–Cas7.4) stack along the crRNA on one side, forming a filament (Figure 1B). It is worth noting that residues 1031–1390 (Insertion domain) in the Cas7.4 domain were not observed in the cryo-EM maps due to the flexibility. The Cas7 domains in gRAMP display RRM folds similar to that observed in type III-A Csm3 and III-B Cmr4, which are composed of a four-stranded antiparallel β-sheet flanked by three α-helices on one side. A significant difference between the Cas7 domain in gRAMP and Csm3/Cmr4 is that each of the Cas7 domain contains a zinc-finger motif. In the Cas7.1, Cas7.2, and Cas7.4 domains, the zinc ions are coordinated by the C2C2 zinc-finger motif. In contrast, the zinc ion in the Cas7.3 domain is coordinated by the CCCH zinc-finger motif. The Cas11 domain locates on the other side of the crRNA, interacting with Cas7.2, Cas7.3 and target ssRNA (Figure 1B). Similar to the structure of type III-A Csm2 and III-B Cmr5, the Cas11 domain in gRAMP displays a helix bundle conformation, with three α-helices on one side and two α-helices on the other side (Figure 1B). The architecture of type III-E effector system is significantly different from the architecture of type III-A and type III-B. The overall structure of the Cmr/Csm complex displays a capsule-like architecture^9^. Four Cmr4 (three Csm3) and three Cmr5 (two Csm2) subunits constitute the double helical backbone, which is capped by Cmr1–Cmr6 (Csm5) at one end (head) and Cmr2–Cmr3 (Csm1–Csm4) at the other (tail).

The four Cas7 domains and the single Cas11 domain look like a palm to grasp the crRNA: target ssRNA heteroduplex. In the constructed structure, the crRNA is composed of a 19-nt 5’-handle and an 18-nt guide segment. The 5’-handle region (G(−19)-C(−1)) lies in the groove formed by the Cas7.1 and Cas7.2 domains (Figure 1B). The crRNA: target ssRNA heteroduplex (C(1):G(−1)-G(16):C(−16)) is enclosed by the Cas7.2, Cas7.3, and Cas11 domains (Figure 1B). Due to the insertion of thumb-like β-hairpins in the Cas7.2 and Cas7.3 domains, bases in the crRNA: target ssRNA heteroduplex at the fourth and 10th positions flipped outside. Furthermore, a loop (residues 1458–1464) in Cas7.4 passes through the heteroduplex, twisting the bases between positions 13 and 14. The interaction between gRAMP and the crRNA: target ssRNA heteroduplex is mediated by extensive hydrogen bonds and hydrophobic interactions (Figure 1C). As observed in the structure, the C(−16) in the handle region of crRNA forms hydrogen bonds with Lys47 and Lys55 in Cas7.1 (Figure 1C). In Cas7.1, U(−15) forms π-π stacking and hydrogen bonds with Phe57 and Asp157, respectively. U(−14)-A(−12) makes hydrophobic stacking with Arg64, His143, Tyr149, and His152 in Cas7.1. U(−11)-G(−10) forms hydrogen bonds with Lys141 and Lys101 in Cas7.1. U(−9) forms π-π stacking with Phe104 in Cas7.1. C(−8) and G(−5) form hydrogen bonds with Arg510 (Cas7.2) and Lys107 (Cas7.1), respectively. A(−7) forms cation-π stacking with Arg37, and C(−6) forms cation-π stacking with Lys229 and Arg472. G(−4) forms π-π stacking and hydrogen bond with Phe192 and Arg37 in Cas7.1, respectively. G(−3) forms hydrogen bond with Arg481 in Cas7.2. The interactions between gRAMP and the crRNA: target ssRNA heteroduplex are mainly mediated by the sugar-phosphate backbone and the neighboring basic amino acids, including His327, His328, Arg294, Arg323, Lys371, Asp698, Arg728, Arg732, Arg1004, Arg1008, Lys1426, Lys1479, Arg1505, and Lys1553 (Figure 1C). A previous study has shown that gRAMP cleaves target ssRNA at two defined positions, which are located after the third and ninth nucleotides, six nucleotides apart^7^. In the structure of gRAMP-crRNA-ssRNA-TPR-CHAT, residue Arg294 in Cas7.2 neighbors the phosphodiester bonds between the third and fourth nucleotides, whereas residue Asp698 in Cas7.3 neighbors the phosphodiester bonds between nucleotides 9 and 10. To investigate whether the residue Arg294 and Asp698 are responsible for cleaving the target ssRNA, R294A and D698A mutants were constructed. The enzymatic activity results showed that the R294A and D698A mutants abolished the ability to cut target ssRNA in each cleavage site, indicating that residues of Arg294 in Cas7.2 and Asp698 in Cas7.3 are vital for ribonuclease activity (Figure 1D). This study further performed *in vitro* target ssRNA cleavage by the gRAMP using mismatched RNAs. As observed in Figure 1E, mismatches 1/2, 3/4, and 5/6 abolished the target RNA cleavage at site 1. In contrast, mismatch 11/12 abolished the target RNA cleavage at site 2. Mismatch 7/8 decreased the target RNA cleavage at site 2, while mismatch 9/10 decreased the target RNA cleavage at both site 1 and site 2. Interestingly, the cleavage of mismatched RNAs 13/14, 15/16, and 17/18 was improved (Figure 1E,S4). The data suggested that the base pairings surrounding position 3 and 9 are vital for the target RNA cleavage. As you known, the gRAMP and crRNA both contribute to the binding of target RNA. The mismatch between crRNA and tgRNA will seriously weaken the interaction between gRAMP-crRNA and target RNA. So, the product RNA will be more prone to dissociating from the crRNA. The mismatch between crRNA and tgRNA will promote the substrate turnover, thus increasing the endoribonuclease activity.

TPR-CHAT is a 716-residue protein that consists of two domains (Figure 1A). The N-terminal TPR domain displays a 16 α-helix bundle conformation. The C-terminal CHAT domain contains an 11-stranded antiparallel β-sheet flanked by four α-helices on one side and five α-helices on the other side (Figure 1B). Structural alignment using the DALI server indicates that TPR-CHAT exhibit a peptidase fold. TPR-CHAT are located adjacent to Cas7.1, Cas7.3, and the 3’-end of the target ssRNA (Figure 1B). The interface between TPR-CHAT and gRAMP-crRNA-ssRNA buries a solvent-accessible surface area of ∼1435 Å^2^, and is composed of extensive hydrogen bonds (including Lys57, Glu102, Glu438, Asn446, and Gln444 in TPR-CHAT, and Ile452, Ser449, Phe402, Thr406, and Asn502 in gRAMP), salt bridges (including Lys42, Lys43, and Lys333 in TPR-CHAT, and Asp749, Glu745, and Asp184 in gRAMP), and hydrophobic interactions (including Leu449 and Tyr450 in TPR-CHAT, and Val405, Pro407, Pro408, Ile411, His566, Ile507, and Arg511 in gRAMP) (Figure 1B). Notably, TPR-CHAT also makes direct contacts with the target ssRNA (Figure 1C). Residues Lys42, Lys85, and Lys92 form hydrogen bonds with phosphate moieties of A(−3)-C(−4). Residue Asn46 forms hydrogen bond with phosphate moieties of G(−2)-A(−3). Residue Lys50 forms hydrogen bond with phosphate moieties of G(−1)-G(−2) (Figure 1C).

Structural superimposition of gRAMP-crRNA-TPR-CHAT with or without target ssRNA yields a root mean squared deviation of 1.042 Å over 1909 aligned Cα atoms, indicating that target ssRNA binding does not cause the extensive rearrangement of gRAMP and TPR-CHAT (Figure 1F). However, it is worth noting that, due to the direct interaction between the N-terminal TPR domain and the 3’-end of target ssRNA, the TPR domain drew slightly closer to the crRNA: target ssRNA heteroduplex (Figure 1F). TPR-CHAT could enforce the interactions between gRAMP and target ssRNA, which would hinder the dissociation of products from the crRNA and weaken the enzymatic activity. To explore the effect of TPR-CHAT on the target ssRNA cleavage by gRAMP, this study performed an *in vitro* target ssRNA cleavage assay by adding TPR-CHAT of increasing molar concentrations. As observed in Figure 1G, the cleavage activity of gRAMP at site 1 was decreased when the higher concentrations of TPR-CHAT were included in the reaction, further supporting the structural observation.

## Discussion

Taken together, these results provide the atomic view of the architecture of Craspase. Combing mutagenesis experiments and structural information enabled the further identification of two catalytic residues in gRAMP for target ssRNA cleavage. Interestingly, the Insertion domain (residues 1031–1390) was not observed in our structure. It is worth to note that we also try the focused 3D classification and local refinement, however the density of the INS domain is still not observed. We also get a low-resolution model of DiCas7-11 in complex with crRNA and target ssRNA, and the Insertion domain is clearly observed. According to our biochemical experiment, the size of crRNA is approximate 37 bp, which is consistent with our structural observation. So, we speculate that the missing of the density of the Insertion domain is caused by the processing of pre-crRNA. The mismatched RNAs (mismatches 13/14, 15/16, and 17/18) increase the enzymatic activity, possibly due to the mismatched target RNAs (13/14, 15/16, and 17/18) are more prone to dissociating from the crRNA. During the preparation of this manuscript, the structure of Desulfonema ishimotonii Cas7-11 (DiCas7-11) in complex with crRNA and target ssRNA was reported at 2.5 Å^10^. Sb-gRAMP possesses 34% sequence identity with DiCas7-11 (Figure S5). Structural comparison of these two structures indicates that the overall structure of Sb-gRAMP adopts a similar fold to that of DiCas7-11, with both containing one Cas11 domain and four Cas7 domains (Figure S6). In addition, the crRNA and target ssRNA also display similar conformation. The root mean square deviation (RMSD) of these two structures is 2.05 for the aligned 848 Cα atoms. The major difference between these two structure is that the Cas11 domain rotates about 30 °, and the Insertion domain is not observed in our structure due to flexibility (Figure S6). Recently, Nishimasu’s group has resolved the cryo-EM structures of the DiCas7-11-crRNA-Csx29 complex with and without target RNA^11^. In the structure of DiCas7-11-crRNA-Csx29-tgRNA, the CTD of Csx29 (residues 75-751) is not observed. The author proposed that upon the tgRNA binding, Csx29 adopts a flexible conformation, which allows the active center to access to the substrate protein. However, in our structure, the density of TPR-CHAT is clearly observed. Structural superimposition of TPR-CHAT with or without target ssRNA yields a root mean squared deviation of 0.72 Å over 637 aligned Cα atoms, indicating that target ssRNA binding does not cause the extensive rearrangement of TPR-CHAT, whereas the helix bundle in the TPR domain moves slightly toward crRNA. Thus, our structural observation indicates that the binding of target ssRNA makes the slight conformational changes in the NTD of TPR-CHAT, although the conformation of the CTD of TPR-CHAT remains unchanged. In addition, the protease active center is solvent exposed in the target ssRNA to allow the access of the substrate protein (Figure S7). We just noticed that Ke’s group have resolved the structures of Craspase in four different states, including resting state, non-matching PFS RNA bound state, matching PFS RNA bound state, and non-matching PFS RNA post-cleavage state. They found that the binding of target RNA with a non-matching PFS will cause a rigid-body movement of CHAT domain, which results the distance of catalytic residues C627 and H585 from 6.6 to 3.3 Å. However, we didn’t observe the dramatic conformational changes in CHAT domain. This may be due to the length of PFS sequence observed in our structure is 3 bp shorter than that is resolved by Ke’s group. Previous study supports the possibility that the III-E CRISPR–Cas system could trigger abortive infection using a caspase-like peptidase, whereby host cells suicide to prevent phages from spreading beyond the infected cell. Nishimasu’s group has revealed that the caspase-like peptidase TPR-CHAT functions to cleave the type III-E associated Csx30 and releases the toxic N-terminal fragment, which in turn triggers the cell growth arrest or cell death through inhibiting the RpoE activity^12^.

In summary, the data presented in this manuscript reveal the mechanism of recognition of crRNA and target ssRNA by gRAMP. The structures constructed here provide insights into pre-crRNA processing and target ssRNA cutting. Furthermore, this work reports the structure of the CRISPR type III-E effector in complex with the binding partner TPR-CHAT, which provides vital clues for elucidating the functional relation between the CRISPR-Cas system and caspase peptidase.

## Materials and Methods

### Strains and plasmids

The full-length genes of gRAMP and crRNA were synthesized by GENEWIZ (Azenta Life Sciences), subcloned into the bacterial expression vector pACYCDuet-1 MCS I and MCS II (An N-terminal 6 His-Sumo-Twin-Strep-Tag was added in pACYCDuet-1). The cDNAs of full-length TPR-CHAT were synthesized using the bacterial expression vector pGEX-6P-1 (GE Healthcare, with an N-terminal GST tag). Point mutations were introduced using Fast Site-Directed Mutagenesis Kit in the *Sb*-gRAMP gene and verified by DNA sequencing. Primers used in the mutation research were listed in Table S1. The GenBank ID of gRAMP is KHE91659.1. The GenBank ID of TPR-CHAT is KHE91663.1. The DNA sequence of crRNA was listed in Table S2. The DNA sequences of the the Spacer and Repeat in the Candidatus “Scalindua brodae” CRISPR array were listed in Table S3.

### Protein expression and purification

The TPR-CHAT plasmid was co-transformed with the plasmid of *Sb*-gRAMP & crRNA into *E. coli* Bl21 using ampicillin and chloramphenicol resistance. Expression of the recombinant protein, *Sb*-gRAMP-crRNA-TPR-CHAT, was induced by 0.3 mM isopropyl β-D-1-thiogalactopyranoside (IPTG) at 20°C. After overnight induction, the cells were collected by centrifugation, and cells were resuspended in buffer A (25 mM Tris·HCl, pH 8.0, 150 mM NaCl) supplemented with 2 mM protease-inhibitor PMSF (phenyl methane sulphonyl fluoride, Sigma). The cells were subjected to lysis by sonication, and cell debris was removed by centrifugation at 23,708 g for 40 min at 4°C. The lysate was first purified using Streptactin beads. The beads were washed, and the bound complexes were cleaved by precision protease in buffer A overnight at 4°C to remove the 6 His-Sumo-Twin-Strep-tag and GST tag. The cleaved complexes were eluted from Streptactin beads and further fractionated by a HiTrap heparin column (AKTA Pure, GE Healthcare) in buffer B (25 mM Tris·HCl, pH 8.0, 3 mM DTT) and buffer C (25 mM Tris·HCl, pH 8.0, 1 M NaCl, 3 mM DTT). Then, the complex was incubated with the ssRNA at a molar ratio of 1: 0.8 at 4°C for 1 h. The complex was applied onto superdex200 increase 10/300 GL column via fast protein liquid chromatography (AKTA Pure, GE Healthcare) with buffer A. The processes of expression and purification of the protein, TPR-CHAT, were the same as the methods of *Sb*-gRAMP-crRNA-TPR-CHAT complex. The cleaved TPR-CHAT protein was eluted from GST beads and further fractionated by a Resource Q (AKTA Pure, GE Healthcare) in buffer. B (25 mM Tris·HCl, pH 8.0, 3 mM DTT) and buffer C (25 mM Tris·HCl, pH 8.0, 1 M NaCl, 3 mM DTT).

### In vitro transcription and purification of ssRNA

The ssRNA was transcribed in vitro using T7 polymerase and purified using 10% concentration denaturing polyacrylamide gel electrophoresis. Transcription template (dsDNA) for ssRNA was generated by polymerase chain reaction (PCR). The reactions were processed at 37°C for 5 h in buffer containing 0.1 M HEPES·K (pH 7.9), 12 mM MgCl_2_, 30 mM DTT, 2 mM spermidine, 2 mM of each NTP, 160 μg ml^− 1^ home-made T7 polymerase, and 500 nM transcription template. The reactions were stopped by letting the samples stand for 1 h at − 80°C. Pyrophosphate was precipitated with Mg^2+^ at cold temperatures, and DNA templates were precipitated with spermidine. The precipitate was removed by centrifugation at 2,514 g for 40 min at 4°C. The RNA was precipitated by 3 M sodium acetate and ethanol overnight. The RNA containing pellets was then resuspended and purified by gel electrophoresis on a denaturing (8 M urea) polyacrylamide gel. RNA bands were excised from the gel and recovered with an Elutrap System followed by ethanol precipitation. RNA was resuspended in diethyl pyrocarbonate H_2_O and stored at − 80°C.

### Cleavage activity assays

Cleavage assays were performed in a 10 μL reaction (25 mM Tris·HCl, pH 8.0, 40 mM NaCl, 2 mM MgCl_2_, and one unit RNase inhibitor) containing 600 nM *sb*-gRAMP-crRNA complex and 166 nM 5’ Cy7-labeled RNA substrate. The *sb*-gRAMP-crRNA complex was first added, followed by the 5’ Cy7-labeled RNA substrate. When exploring the effect of TPR-CHAT during the target RNA cleavage procession of gRAMP, the TPR-CHAT was added before the addition of 5’ Cy7-labeled RNA substrate. Cleavage reactions were conducted at 37°C for 45 min. Reactions were stopped by adding 2× TBE (tris-borate-EDTA)-urea loading buffer. Cleavage products were run on 10% TBE-urea denaturing gel at room temperature in 1× TBE running buffer and visualized by fluorescence imaging. The full match and mismatched RNA (5’ Cy7-labeled) squences used in the cleavage assay were listed in Table S4.

### Cryo-EM data collection

For all of the samples, 3 μL of *Sb*-gRAMP-crRNA-TPR-CHAT and *Sb*-gRAMP-crRNA-ssRNA-TPR-CHAT at 6.0 mg/mL were applied onto glow-discharged 300-mesh R1.2/1.3 Quantifoil grids. The grids were blotted for 3 s and rapidly cryocooled in liquid ethane using a Vitrobot Mark IV (Thermo Fisher Scientific) at RT and 100% humidity. The samples were imaged in a Titan Krios cryo-electron microscope (FEI) with a GIF energy filter (Gatan) at a magnification of 130,000× (corresponding to a calibrated sampling of 1.1 Å per pixel). Micrographs were recorded by EPU software (Thermo Fisher Scientific) with a Gatan K2 Summit direct electron detector, where each image was composed of 40 individual frames with an exposure time of 7.5 s and a dose rate of 8.0 electrons per second per Å^2^. A total of 5,472 (*Sb*-gRAMP-crRNA-TPR-CHAT) and 1,923 (*Sb*-gRAMP-crRNA-ssRNA-TPR-CHAT) movie stacks were collected with a defocus range of − 1.5–− 2.5 μm.

### Image processing

For the structures of *Sb*-gRAMP-crRNA-TPR-CHAT and *Sb*-gRAMP-crRNA-ssRNA-TPR-CHAT, a total of 5,472 and 1,923 micrographs were obtained for data processing, respectively (Extended Data Fig. 1–2). All of the micrographs were motion-corrected using MotionCor2^13^, and the contrast transfer function (CTF) was determined using CTFFIND4^14^. All of the particles were autopicked using RELION3.1^15^ and further checked manually, yielding 4,705,908 and 1,856,129 particles for *Sb*-gRAMP-crRNA-TPR-CHAT and *Sb*-gRAMP-crRNA-ssRNA-TPR-CHAT, respectively. Then, two rounds of two-dimensional classification were performed in RELION3.1 to remove poor two-dimensional class averages. In total, 1,430,971 and 1,351,569 particles from the best classes were selected from *Sb*-gRAMP-crRNA-TPR-CHAT and *Sb*-gRAMP-crRNA-ssRNA-TPR-CHAT, respectively, and they were used for three-dimensional (3D) classification in RELION3.1 to remove poor 3D classes. Classes that showed clear secondary structure features were selected, and they were re-extracted without binning (using a pixel size of 1.1 Å) in RELION3.1. Then, the particles were transferred to cryoSPARC for another round of heterogeneous refinement. For *Sb*-gRAMP-crRNA-TPR-CHAT and *Sb*-gRAMP-crRNA-ssRNA-TPR-CHAT, 114,630 and 87,226 particles from the best class were subjected to non-uniform refinement, respectively. Finally, a 2.9 Å map for *Sb*-gRAMP-crRNA-TPR-CHAT and a 3.0 Å map for *Sb*-gRAMP-crRNA-ssRNA-TPR-CHAT were obtained. The resolutions for the final maps were estimated based on the 0.143 criterion of the FSC curve^16^.

### Model building

Model building was performed based on the 2.9 Å and 3.0 Å reconstruction maps. The TPR-CHAT, gRAMP, crRNA, and target ssRNA were all built de novo using COOT^17^. The two models were refined using the phenix.real_space_refine^18^ application with secondary structure and geometry restraints. The final models were evaluated by MolProbity^19^ and Ramachandran plot^20^. The statistics of the map reconstruction and model refinement are presented in Table S5.

## DATA AVAILABILITY

The atomic coordinates of the gRAMP-crRNA-TPR-CHAT and gRAMP-crRNA-ssRNA-TPR-CHAT complexes have been deposited to the Protein Data Bank under the accession numbers 7Y8T and 7Y8Y, respectively. The corresponding maps have been deposited in the Electron Microscopy Data Bank under the accession numbers EMD-33685 and EMD-33686, respectively.

## ACKNOWLEDGEMENTS

We thank Anqi Zhang and Changyou Guo of the EM platform at School of Life Science and Technology of Harbin Institute of Technology for sample screening and data collection. This research was funded by the National Natural Science Foundation of China (grant no. 31825008 and 31422014), the Tencent Foundation, and Heilongjiang Touyan Team HITTY-20190034 to Z.W.H., and the fellowship of China National Postdoctoral Program for Innovative Talents (grant no. BX20200110) to M.H.G..

## AUTHOR CONTRIBUTIONS

S.W., M.G. and Z.L. prepared the samples and performed functional assays; S.W. and M.G. performed negative staining; A.Z. and C.G. prepared the cryo-EM grids and performed cryo-EM data collection. Y.Z. and Z.H. performed cryo-EM data processing and model building; S.W., Y.Z., and Z.H. wrote the manuscript; Z.H. supervised the study and edited the manuscript.

## Figure legends

**Fig. S1.**
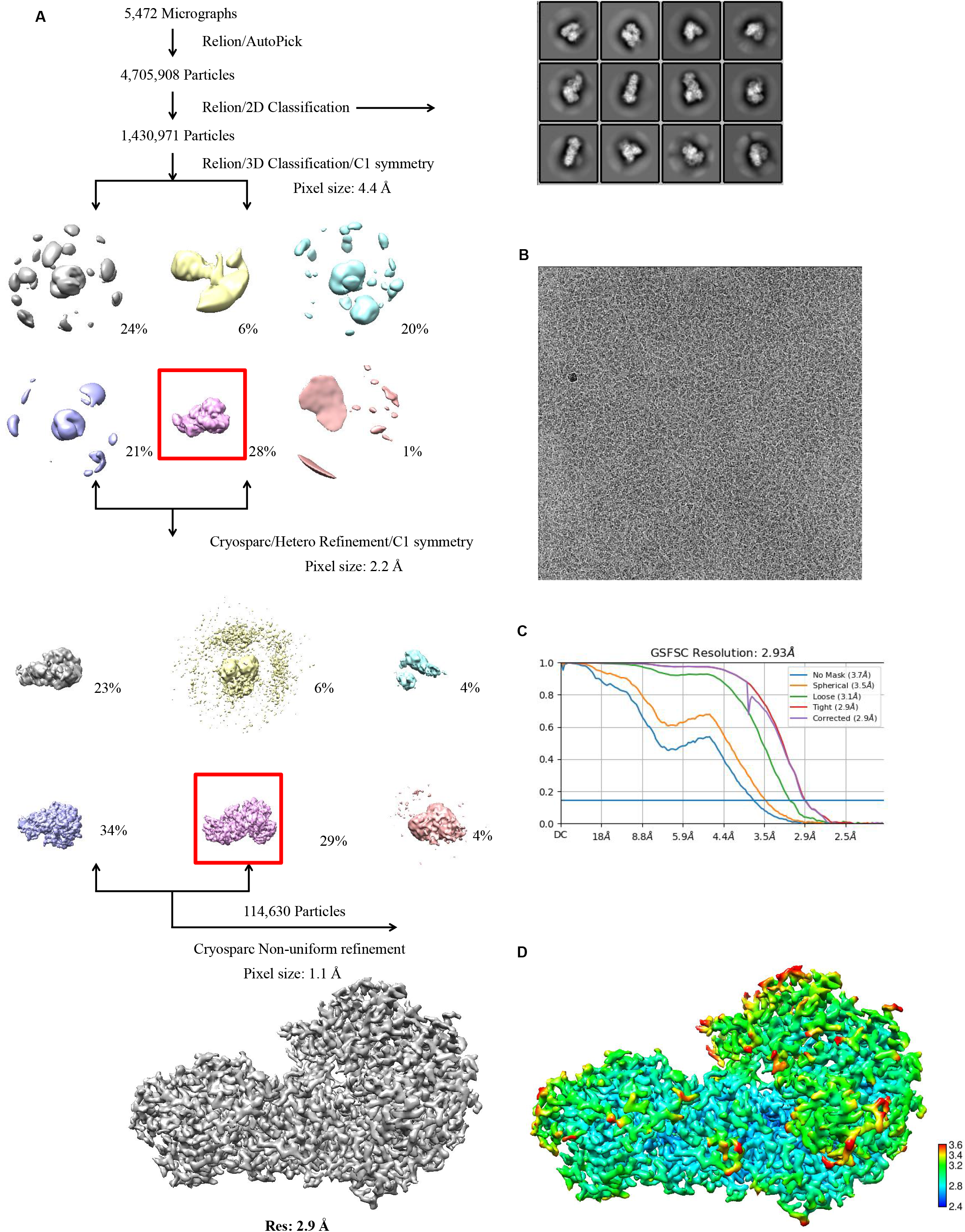
Cryo-EM image processing procedure of the *Sb*-gRAMP-crRNA-TPR-CHAT complex. (A) Image processing workflow of the *Sb*-gRAMP-crRNA-TPR-CHAT complex. (B) A representative raw cryo-EM image of the *Sb*-gRAMP-crRNA-TPR-CHAT complex. (C) Gold-standard FSC plot for the final three-dimensional (3D) reconstruction of the *Sb*-gRAMP-crRNA-TPR-CHAT complex. (D) Resolution map for the final 3D reconstruction of the *Sb*-gRAMP-crRNA-TPR-CHAT complex.

**Fig. S2.**
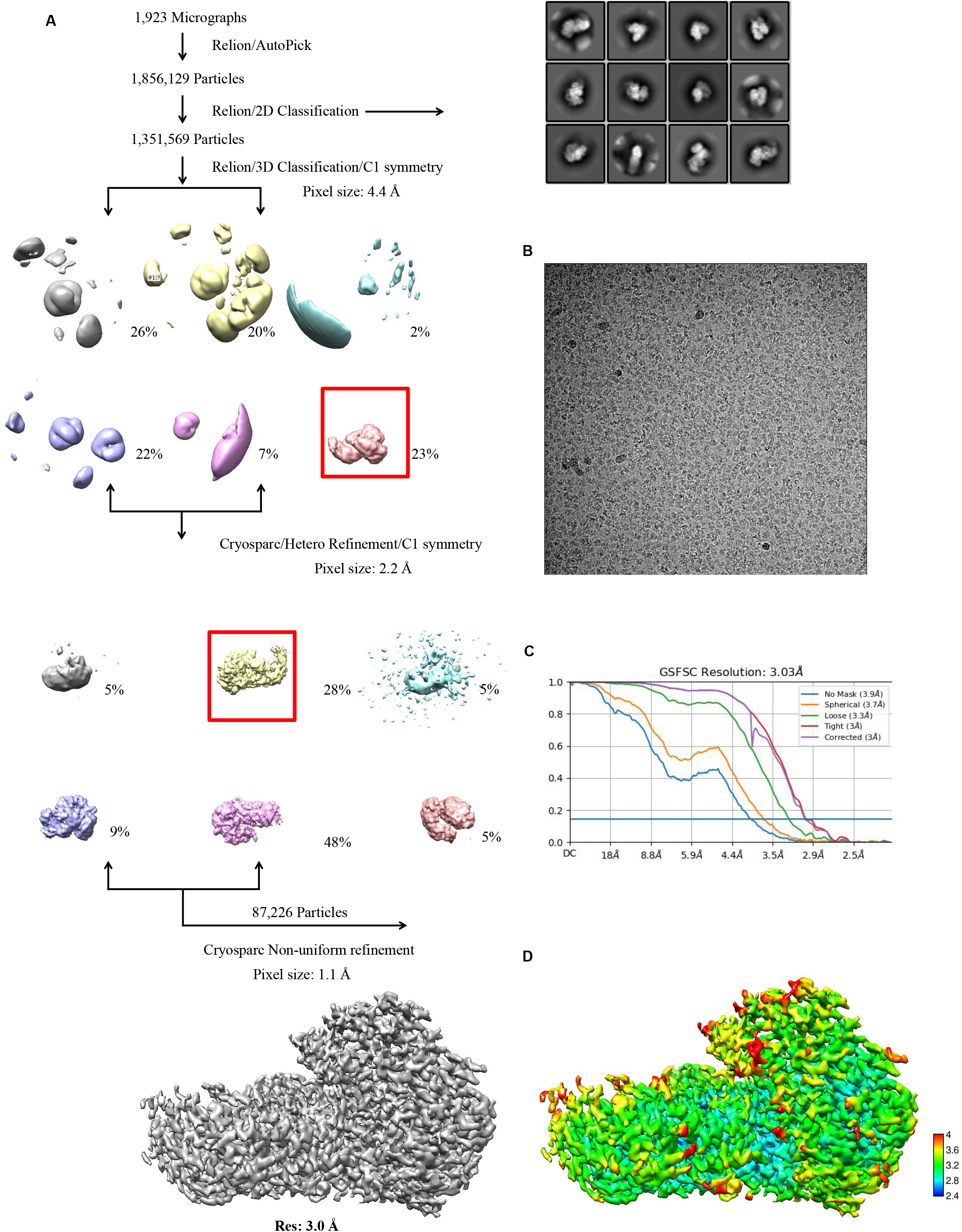
Cryo-EM image processing procedure of the *Sb*-gRAMP-crRNA-ssRNA-TPR-CHAT complex. (A) Image processing workflow of the *Sb*-gRAMP-crRNA-ssRNA-TPR-CHAT complex. (B) A representative raw cryo-EM image of the *Sb*-gRAMP-crRNA-ssRNA-TPR-CHAT complex. (C) Gold-standard FSC plot for the final 3D reconstruction of the *Sb*-gRAMP-crRNA-ssRNA-TPR-CHAT complex. (D) Resolution map for the final three-dimensional (3D) reconstruction of the *Sb*-gRAMP-crRNA-ssRNA-TPR-CHAT complex.

**Fig. S3.**
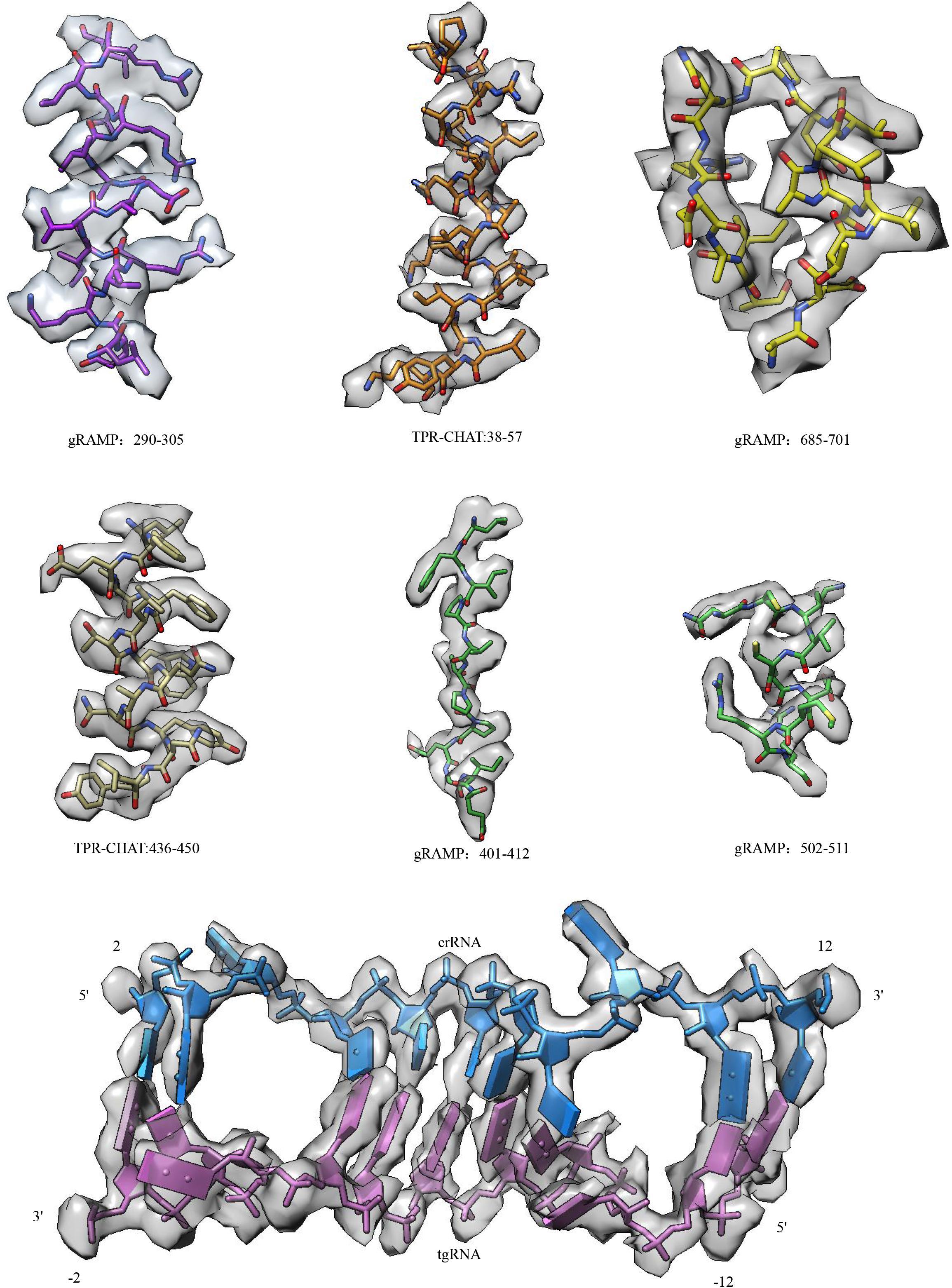
EM densities for indicated regions of the atomic model. EM densities for indicated regions of gRAMP, TPR-CHAT and crRNA: target ssRNA heteroduplex. The color is the same as Figure 1B.

**Fig. S4.**
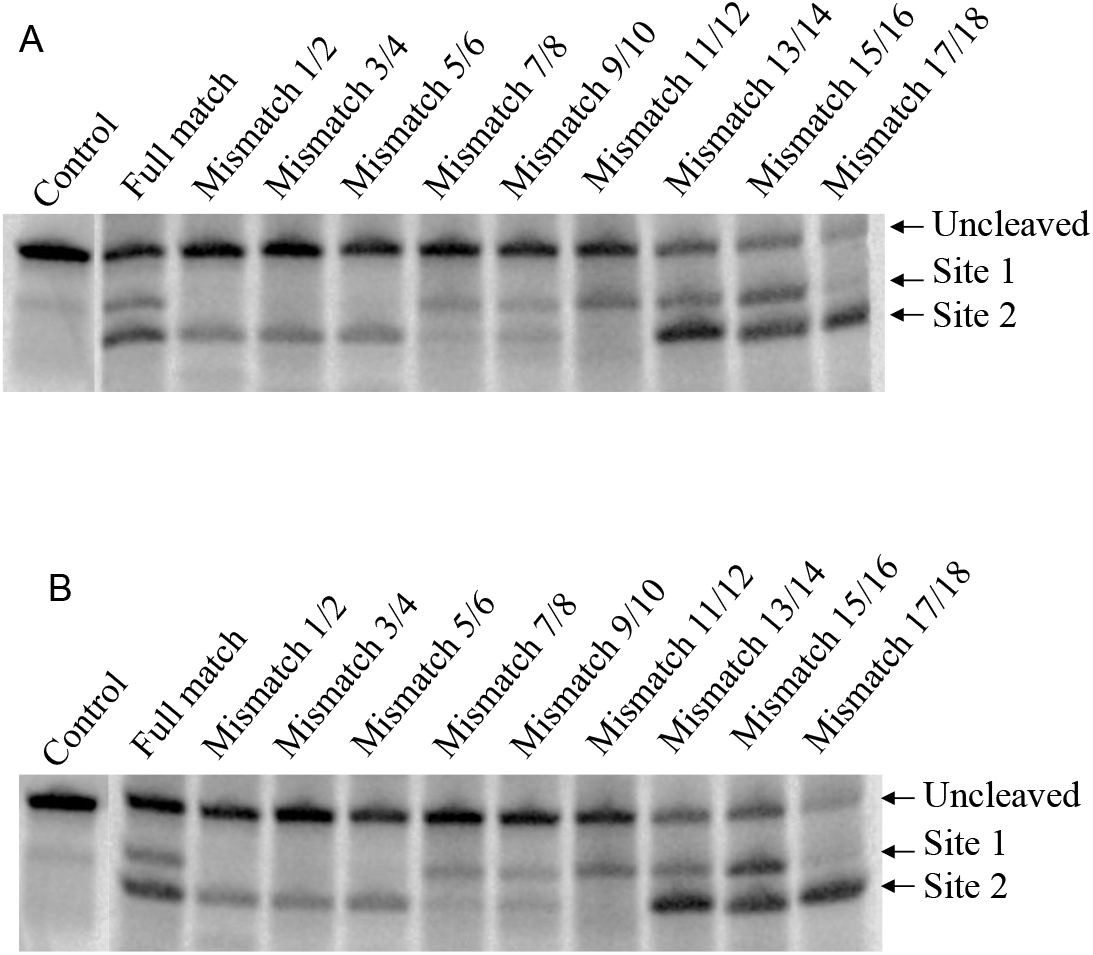
In vitro target ssRNA cleavage by the WT gRAMP using mismatched RNAs. Multiple replicates of Figure 1E were performed (A and B).

**Fig. S5.**
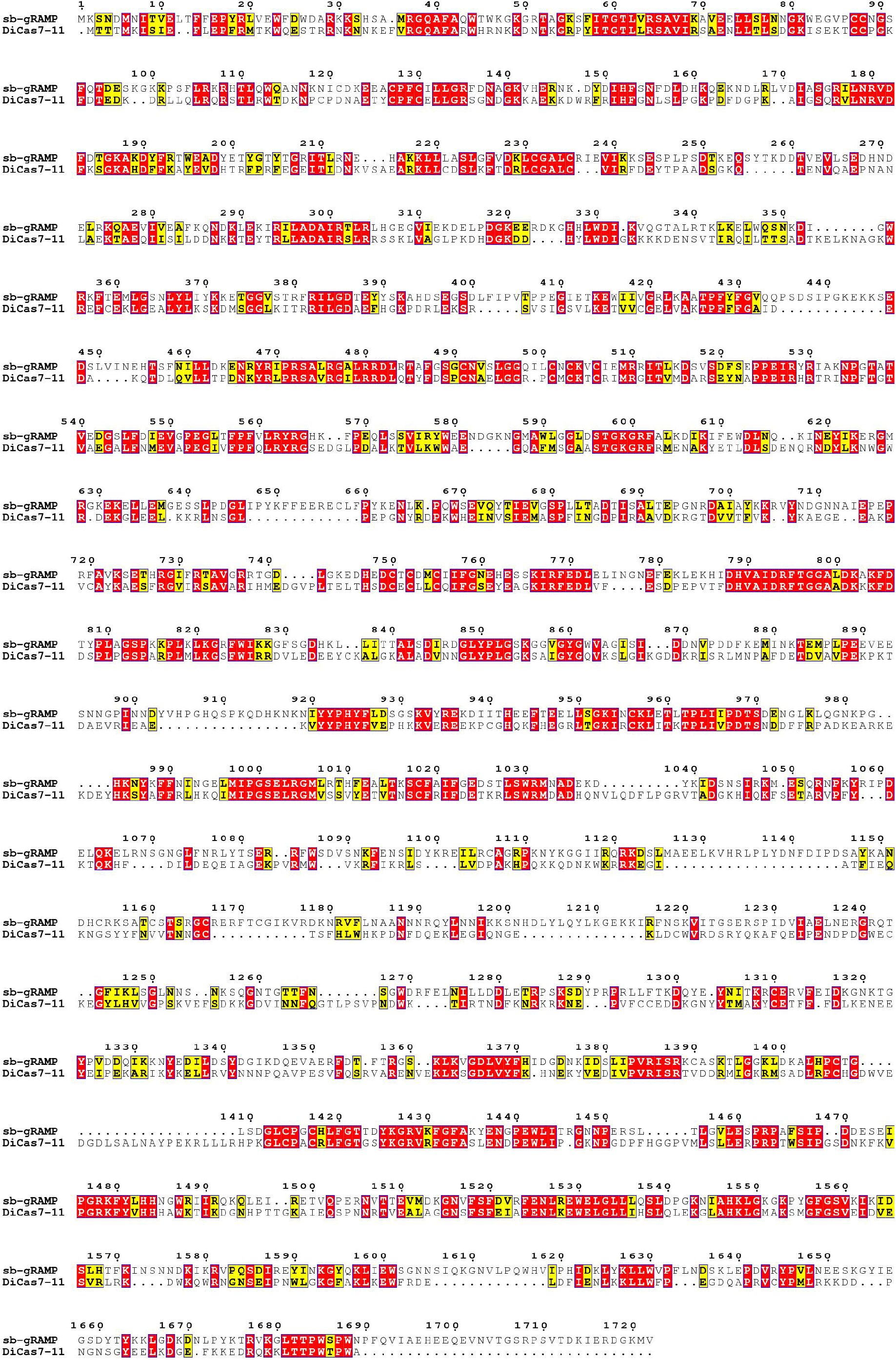
Multiple sequence alignment of the Cas7-11 orthologs. Alignment of *Scalindua brodae* Cas7-11 (*Sb*-gRAMP) and *Desulfonema ishimotonii* Cas7-11. The residues conserved in the Cas7-11 paralogs are highlighted in red.

**Fig. S6.**
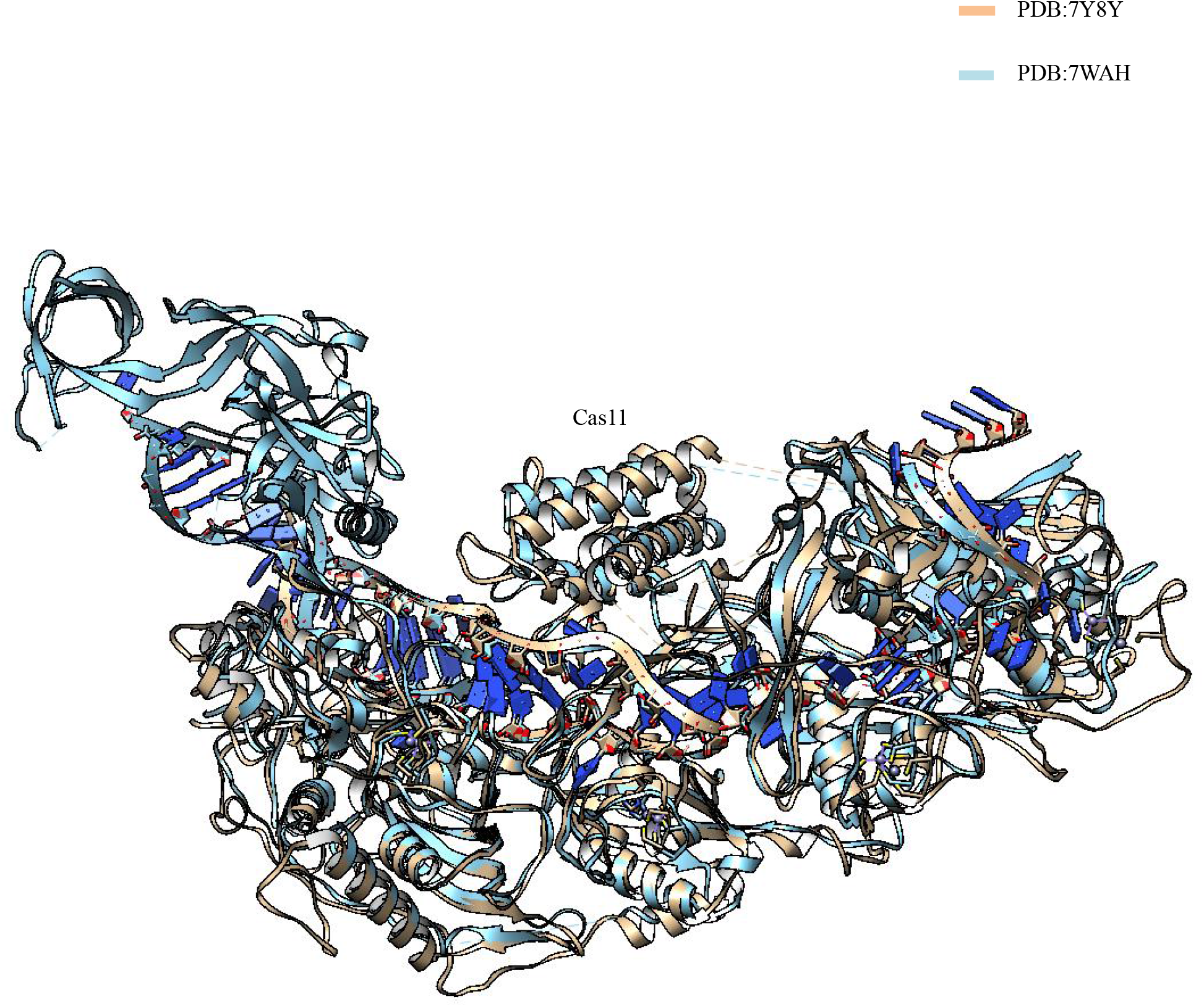
Structural superposition of gRAMP-crRNA-ssRNA and DiCas7-11-crRNA-ssRNA. Structural comparison of gRAMP-crRNA-ssRNA and DiCas7-11-crRNA-ssRNA. The gRAMP-crRNA-ssRNA is shown in orange, and the DiCas7-11-crRNA-ssRNA is shown in cyan.

**Fig. S7.**
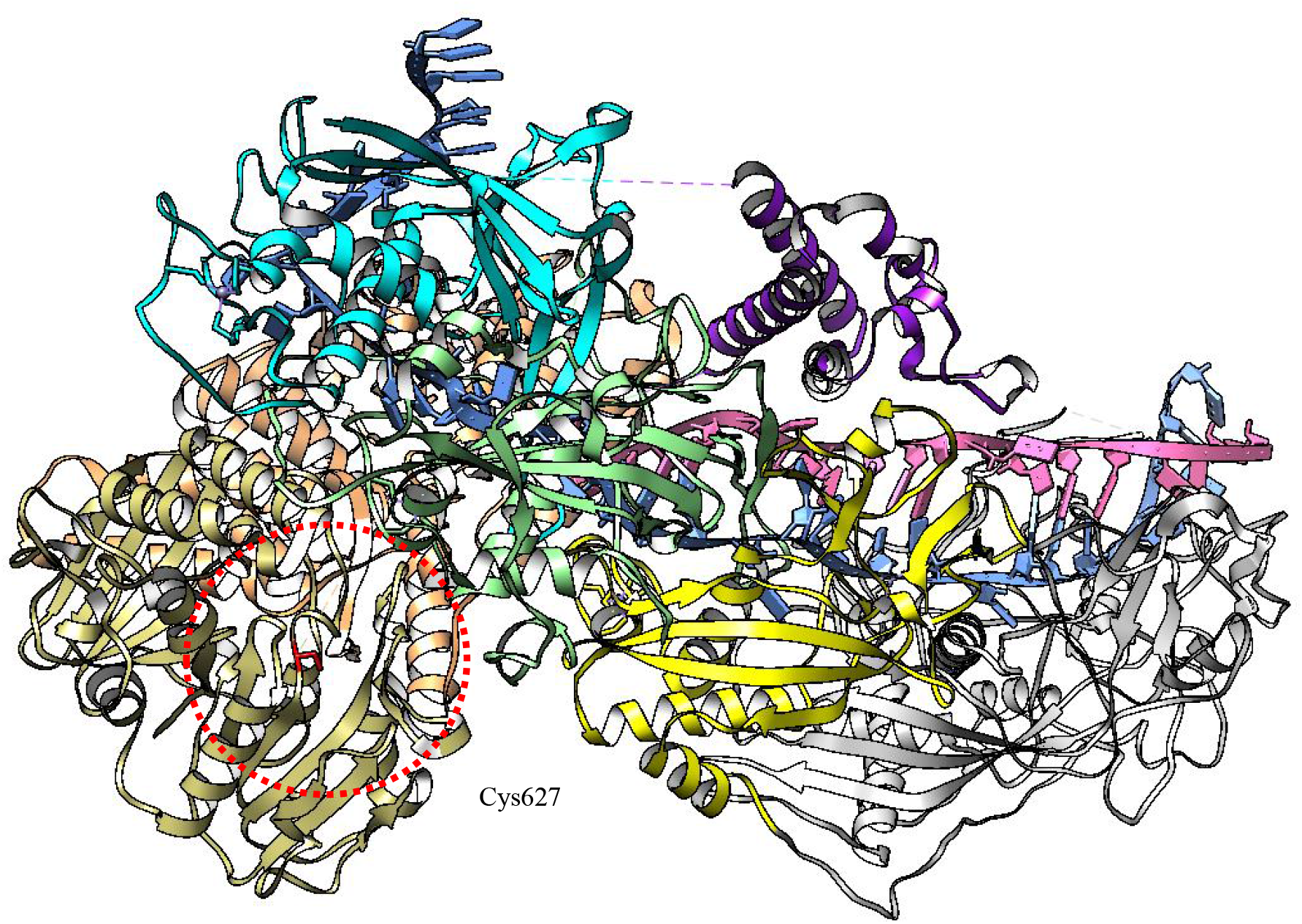
Cartoon show of the structure of gRAMP-crRNA-ssRNA-TPR-CHAT. Overall structure of the gRAMP-crRNA-ssRNA-TPR-CHAT complex. The active center of TPR-CHAT is indicated as red dotted line. The catalytic residue Cys627 is labelled. The color of gRAMP-crRNA-ssRNA-TPR-CHAT complex is the same as Figure 1B.

**Table S1.**
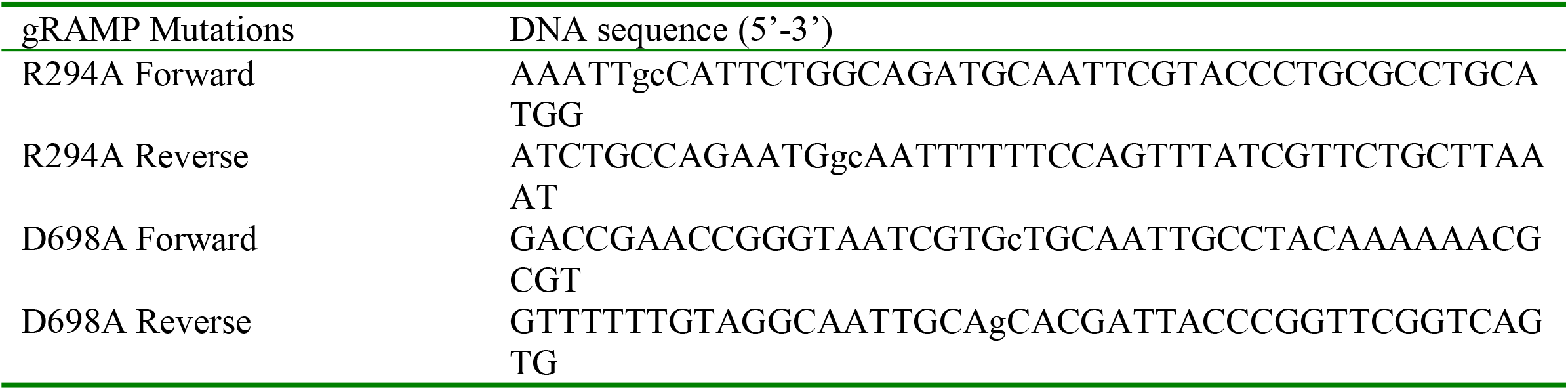
Primers used in the gRAMP mutation research.

**Table S2.**
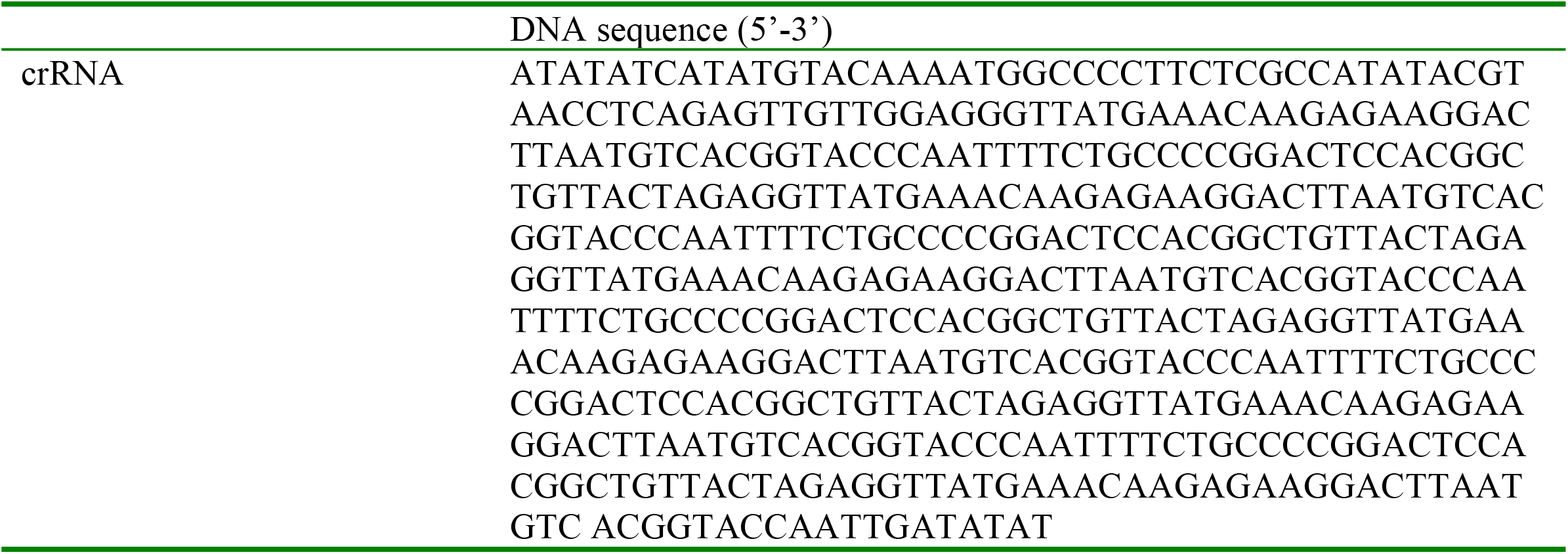
DNA sequence of the crRNA in the Candidatus “Scalindua brodae” CRISPR array.

**Table S3.**
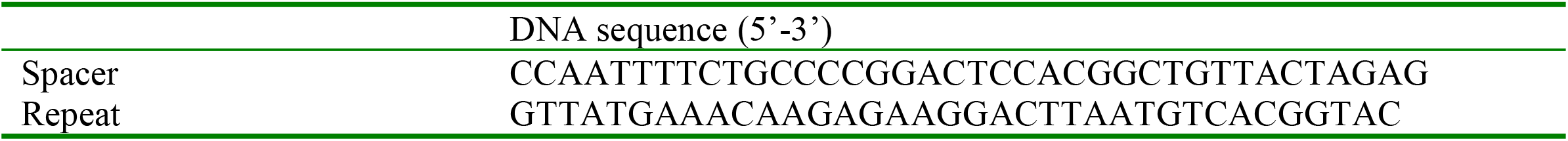
DNA sequences of the Spacer and Repeat in the Candidatus “Scalindua brodae” CRISPR array.

**Table S4.**
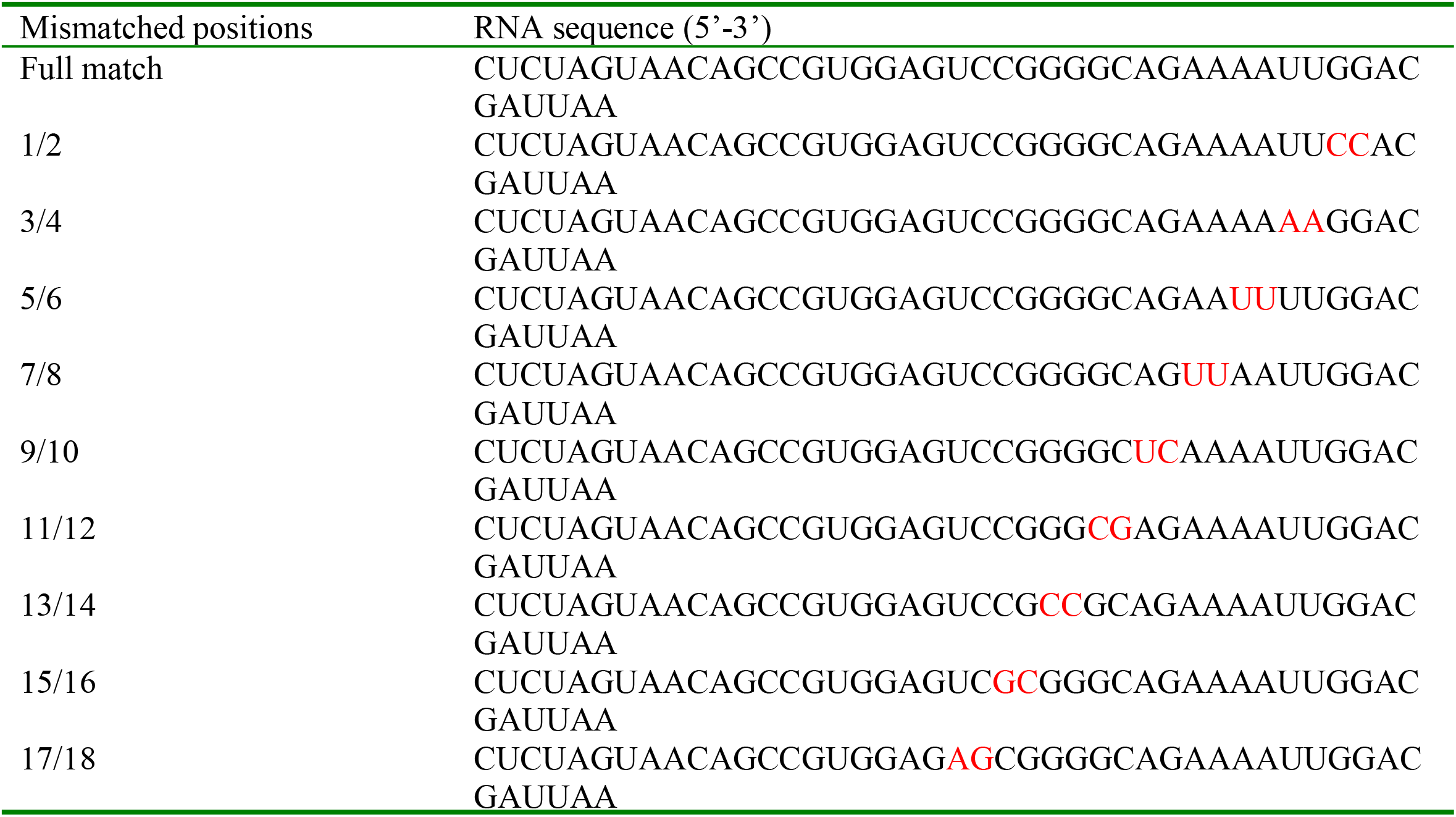
Full match and Mismatched RNA (5’ Cy7-labeled) sequences used in the cleavage assay.

**Table S5.** Cryo-EM data collection, refinement, and validation statistics.

|  | gRAMP-crRNA-TPR-CHAT<br>(EMD-33685)<br>(PDB-7Y8T) | gRAMP-crRNA-ssRNA-TPR-CHAT<br>(EMD-33686)<br>(PDB-7Y8Y) |
| --- | --- | --- |
| **Data collection and processing** |  |  |
| Magnification | 130,000 | 130,000 |
| Voltage (kV) | 300 | 300 |
| Electron exposure (e-/Å<sup>2</sup>) | 60.0 | 60.0 |
| Defocus range (μm) | -1.5~-2.5 | -1.5~-2.5 |
| Pixel size (Å) | 1.1 | 1.1 |
| Symmetry imposed | C1 | C1 |
| Initial particle images (no.) | 396,021 | 313,321 |
| Final particle images (no.) | 114,630 | 87,226 |
| Map resolution (Å) | 2.9 | 3.0 |
| FSC threshold | 0.143 | 0.143 |
| Map resolution range (Å) | 5.0-2.9 | 5.0-3.0 |
| **Refinement** |  |  |
| Initial model used (PDB code) | N/A | N/A |
| Model resolution (Å) | 2.9 | 3.0 |
| FSC threshold | (0.143) | (0.143) |
| Model resolution range (Å) | 5.0-2.9 | 5.0-3.0 |
| Map sharpening *B* factor (Å<sup>2</sup>) | -99.0 | -104.7 |
| Model composition |  |  |
| Nonhydrogen atoms | 16754 | 17023 |
| Protein residues | 1978 | 1971 |
| Nucleotide | 37 | 55 |
| Ligand | 4 | 4 |
| *B* factors (Å<sup>2</sup>) |  |  |
| Protein | 51.48 | 49.71 |
| Nucleotide | 61.08 | 73.18 |
| Ligand | 108.64 | 98.72 |
| R.m.s. deviations |  |  |
| Bond lengths (Å) | 0.004 | 0.003 |
| Bond angles (°) | 0.631 | 0.606 |
| **Validation** |  |  |
| MolProbity score | 1.72 | 2.07 |
| Clashscore | 4.70 | 6.43 |
| Poor rotamers (%) | 1.21 | 2.62 |
| Ramachandran plot |  |  |
| Favored (%) | 93.80 | 94.02 |
| Allowed (%) | 6.00 | 5.83 |
| Disallowed (%) | 0.20 | 0.15 |

